# Molecular interactions of the M and E integral membrane proteins of SARS-CoV-2

**DOI:** 10.1101/2021.04.29.442018

**Authors:** Viviana Monje-Galvan, Gregory A. Voth

## Abstract

Specific lipid-protein interactions are key for cellular processes, and even more so for the replication of pathogens. The COVID-19 pandemic has drastically changed our lives and cause the death of nearly three million people worldwide, as of this writing. SARS-CoV-2 is the virus that causes the disease and has been at the center of scientific research over the past year. Most of the research on the virus is focused on key players during its initial attack and entry into the cellular host; namely the S protein, its glycan shield, and its interactions with the ACE2 receptors of human cells. As cases continue to raise around the globe, and new mutants are identified, there is an urgent need to understand the mechanisms of this virus during different stages of its life cycle. Here, we consider two integral membrane proteins of SARS-CoV-2 known to be important for viral assembly and infectivity. We have used microsecond-long all-atom molecular dynamics to examine the lipid-protein and protein-protein interactions of the membrane (M) and envelope (E) structural proteins of SARS-CoV-2 in a complex membrane model. We contrast the two proposed protein complexes for each of these proteins, and quantify their effect on their local lipid environment. This ongoing work also aims to provide molecular-level understanding of the mechanisms of action of this virus to possibly aid in the design of novel treatments.

## Introduction

Since the beginning of the COVID-19 pandemic, research progress has been unprecedented to elucidate and block the progression of SARS-CoV-2 infection, the virus that causes the disease. Scientists have been collaborating as never before in open platforms, sharing data as soon as available to aid in the fight against the virus. Though other coronaviruses have been around for several years,^1–3^ the rate of transmission, infectivity, and unpredictability of SARS-CoV-2 make for an urgent need to understand its mechanisms at the molecular level. Several companies have provided viable vaccine candidates to the global market to combat the spread of the virus; yet, novel mutants and the rate of transmission clearly show the need for more and improved alternatives.

Viruses have developed mechanisms to use host cell proteins and lipids to propagate infection.^4, 5^ Understanding these interactions at the molecular level – and how they modulate the structure, aggregation, and function of viral proteins – will provide new insights about the mechanisms of viral replication. The design of successful novel therapeutic approaches against SARS-CoV-2 infection depends largely on the knowledge of such mechanisms and identifying the weak points of the viral life cycle.

Like other coronaviruses, SARS-CoV-2 hast four structural envelope proteins: the spike (S), nucleocapsid (N), envelope (E), and membrane or matrix (M).^1^ Of these, S, E, and M have transmembrane regions that are embedded in the lipid envelope that encloses the viral genome, a positive RNA strand. The N protein mainly interacts with the RNA itself, recruiting it to viral assembly sites and interacting with membrane lipids and other structural proteins to incorporate its cargo into new viral particles.^6, 7^ S is the key player during viral infection, interacting with ACE2 receptors on the surface of cells targeted by the virus and undergoing several conformational changes and proteolytic cleavages that enable viral fusion and release of the viral genome into the cellular host.^8^ M has been identified as a main orchestrator of viral assembly, interacting with all the other structural proteins at some point to recruit them and stabilize the formation of new viral particles at the endoplasmic reticulum-Golgi intermediate compartment (ERGIC).^8–10^ The E protein is classified as a viroporin,^11^ working as an ion channel for the viral particle and blocking the immune response from the host during viral replication.^2, 10, 12, 13^ It is still unclear how the E protein is involved in the assembly process and/or scission of newly formed viral particles.^1, 14^

Very few studies are currently available on the role and dynamics of the M and E integral membrane proteins of SARS-CoV-2.^11, 15–17^ These are both known to be key for viral assembly and infectivity,^9, 16^ and the M protein could potentially serve as an antigen.^18^ Latest experimental studies examining the interactions among structural proteins of the virus identified M and E are both required to recruit and retain S proteins at the viral assembly site.^17^ Yet, their precise mechanisms and relevance in the viral life cycle remain largely unclear. By far, the main focus of current research is towards understanding and blocking the interaction of the S proteins on the surface of the virus. Most of the treatments consist of repurposing existing drugs to block different stages of the viral cycle to stop viral replication, and primarily to block the S-ACE2 initial interaction.^19^ Current vaccine candidates also work by eliciting an immune response in the human body such that it produces antibodies that will recognize and bind to the S protein in case of an infection.^18, 20^ Despite great advances against the virus, we are still in urgent need to gain better understanding of the viral life cycle in its entirety to inspire and repurpose drug therapies more efficiently as the virus mutates.

The relevance and involvement of lipids in cellular and viral processes has been well stablished over the past two decades.^4, 5, 21^ In this work, we take a closer look at the interplay between lipids and the M and E proteins of SARS-CoV-2 at early stages of viral assembly. We present the results of long all-atom molecular dynamics (MD) simulations to study molecular interactions in a complex membrane model, built to mimic the cell organelle where viral assembly takes place, the ERGIC.^3, 22, 23^ We discuss protein-protein interactions in detail for a proposed M dimer as well as the E channel formed by five monomers. At the time of this study, only structures from homology modeling studies were publicly available at the MolSSi Covid-19 Hub.^24, 25^ To add to this emerging body of knowledge, we therefore simulated two proposed models for each protein, shown in Figure 1, and characterized the differences in local membrane response based on protein conformation.

**Figure 1.**
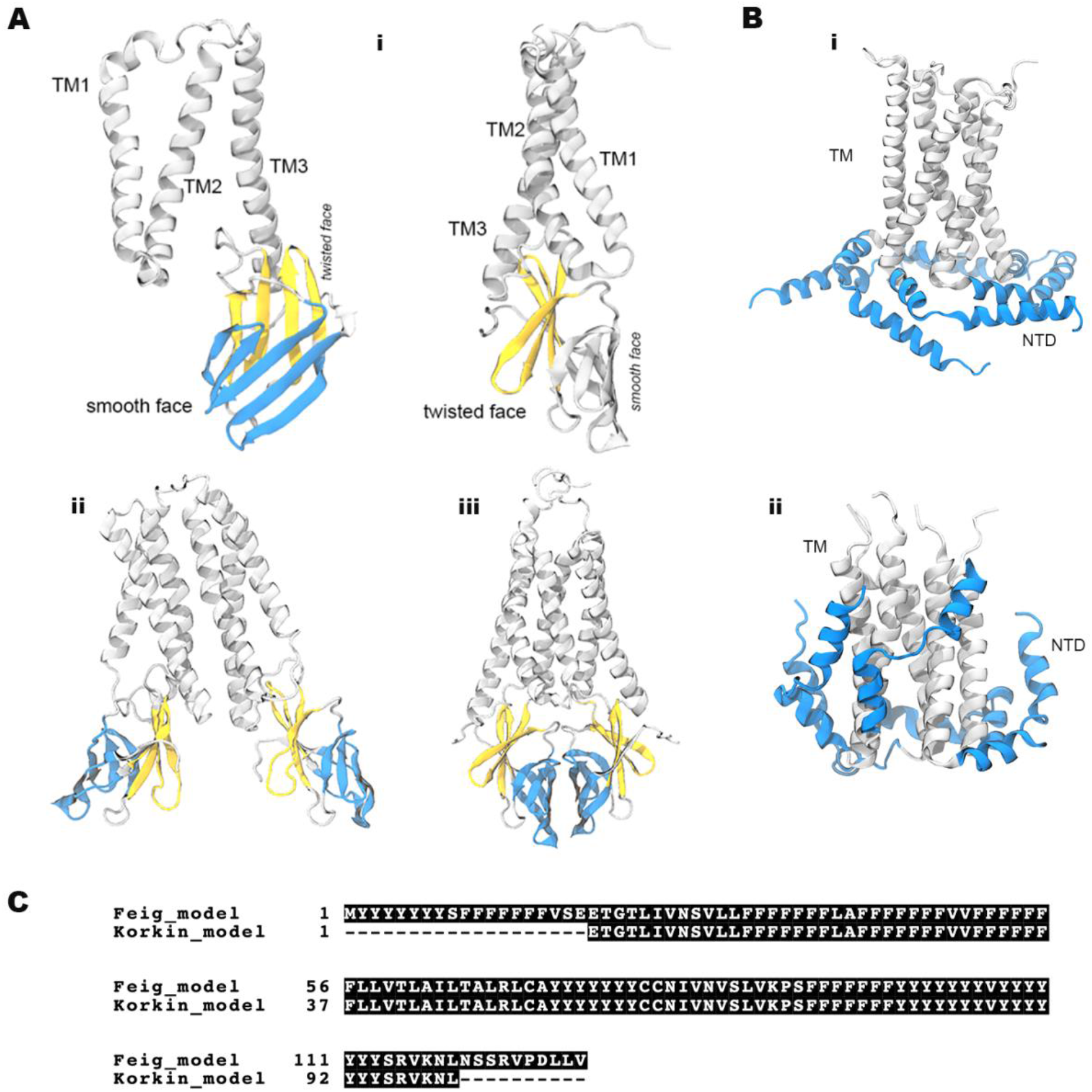
Proposed homology models for integral membrane proteins of SARS-CoV-2. (A) M dimers (Feig lab^23^); (i) Protein regions of an M monomer, three transmembrane helices (TM) and two distinct faces of the C-terminal domain (CTD), which we use to differentiate the (ii) open and (iii) closed conformations. (B) E protein channel from the (i) Feig lab^23^, and (ii) Korkin lab^24^; the transmembrane (TM) helices are shown in grey and the C-terminus helices (CTD) in blue. (C) Sequence alignment between the Feig and Korkin proposed models for the E channel.

As this manuscript was being prepared, two experimental studies were published on the structure and function of the transmembrane region of the E channel,^11, 15^ which we comment on in the discussion section. To our knowledge, there are still no experimental structures for the M protein of SARS-CoV-2 that explicitly examine the macromolecular organization of either of these proteins during viral assembly or in formed viral particles. In light of the most recent studies that clearly identify the high involvement of lipids in the pathogenesis and mechanisms of the virus,^26, 27^ we hope this work lays the groundwork to further examine the complex dynamics between viral proteins and lipids. Specifically, the roles of M and E – both integral membrane proteins of SARS-CoV-2 – during viral assembly, membrane deformation, virus maturation, and reshaping of the viral particle during binding and fusion to a new host. This knowledge is also critical for future advances in treatment and preventive care against Covid-19.

## Methods

There are currently no lipidomic studies on the SARS-CoV-2 virion; we chose to model the endoplasmic reticulum (ER) as our starting point in the assembly pathway of the virus to capture the natural lipid sorting and local rearrangement caused by the proteins. Our membrane model has five representative lipid species for the ER of eukaryotic cells.^28–30^ Incidentally, our lipid composition is nearly identical to that of a model bilayer used in an experimental study carried to elucidate the structure and conformation of the transmembrane region of the E channel of SARS-CoV-2 in an ERGIC-like environment.^11^ That study was published at the time when this manuscript was under preparation. Both the experimental and the simulation membrane models have a mixture of PC, PE, PI, PS, and cholesterol lipids. A very similar composition was also used to model the ER / ERGIC in another computational study published during our data curation. That study is a thorough examination and validation of homology models of the E-channel and its ion channeling function;^14^ their model includes a small percentage of sphingomyelin lipids that our model does not include. Our bilayer model contains DYPC/DYPE/POPI/POPS/Chol in 45:10:12:23:10 molar ratio, with 610 lipids per leaflet over a 19×19 nm surface. This composition was chosen to model the membrane thickness and unsaturation degree of the hydrophobic core in terms of number of double bonds per lipid tail. To our knowledge, this model is the most complex yet to be used in the study of SARS-CoV-2 transmembrane proteins and their dynamics with membrane lipids.

For the M and E proteins, we used structures proposed by the Feig lab from homology modeling and refined computational studies;^24^ the second model for the E protein is based on a known structure for the E channel of SARS, and was proposed by the Korkin lab.^25^ Given the degree of conservation between SARS and SARS-CoV-2,^14^ the homology-derived structures used in this study provide a robust preliminary overview of the interplay between integral membrane proteins of the virus and their local lipid environment. The Feig lab proposed two possible conformations for an M dimer; we differentiate them as the open and closed conformations as shown in Figure 1, depending on the relative orientation of the monomers. The C-terminal domain (CTD) of the proteins points towards the luminal side of the ER or ERGIC, and is located in the interior of formed viral particles or virions. The same group proposed a channel structure formed by five units of the E protein, with the N-terminus domains (NTD) located at the membrane interface (refer to Figure 1). The main difference between that model and the one proposed by the Korkin lab is the location of the NTD; the second structure has these helices inside the bilayer core. As we discuss later in this work, the position of the N-terminus helices results in marked differences in the membrane response. Table 1 summarizes our systems of study; the actual protein coordinates were obtained from the MolSSi Covid-19 Hub (www.covid.molssi.org).

**Table 1.** Simulation systems

| System | Environment | System size (atoms) | Simulation length (ns) | Comp. resource |
| --- | --- | --- | --- | --- |
| open M dimer | water | 137,282 | 400 | Frontera (TACC) |
|  | bilayer<br>(610 lipids/leaflet) | 616,117 | 4000 | Frontera &<br>Anton2 (PSC) |
| closed M dimer | water | 190,643 | 400 | Midway2 (RCC) |
|  | bilayer<br>(610 lipids/leaflet) | 667,837 | 4000 | Frontera &<br>Anton2 |
| E channel – Feig | bilayer<br>(610 lipids/leaflet) | 670,765 | 4000 | Midway2 &<br>Anton2 |
| E channel -<br>Korkin | bilayer<br>(610 lipids/leaflet) | 551,741 | 4000 | Midway2 &<br>Anton2 |

All of our systems where built on CHARMM-GUI *Membrane Builder*, or *Quick-Solvator* for the M dimer in water.^31–35^ Fully hydrated bilayers (40+ water molecules per lipid) were built around the M and E protein complexes, respectively, centering the transmembrane regions of the proteins as closely as possible to the bilayer center. We used KCl salt at 0.15 M concentration to neutralize the systems to run on the isobaric-isothermal ensemble on GROMACS simulation package,^36^ with the CHARMM36m force field.^37^ The protein-water systems were run on Frontera at the TACC, or Midway2 at the RCC of the University of Chicago, as listed in Table 1. Upon initial equilibration on these resources, the production runs of the protein-membrane systems were carried on the Anton 2 machine,^38^ hosted at the Pittsburgh Supercomputing Center (PSC). (See Acknowledgments.)

Initial minimizations of all systems were carried following the 6-step protocol provide on CHARMM-GUI.^39^ Followed by a 16-step restrained protocol over 123 ns to further equilibrate the E channels, to ensure proper relaxation of the lipids around it and prevent it from closing. This setup was adapted from previous studies of the influenza A M2 channel.^40, 41^ Initially, the alpha carbons of the transmembrane helices (TM) of the E channel were restrained and only the N-terminus helices, denoted as alpha-helices (AH) in this work, were allowed to move, followed by the reverse setup and final removal of all restrains prior to starting the production run of these systems. No water molecules were found in the bilayer region during the restrained protocol.

In all our trajectories we used a simulation timestep of 2fs and periodic boundary conditions. The temperature was set to 310.15 K and controlled with the Nose-Hoover thermostat ^42, 43^ with a coupling time constant of 1.0 ps in GROMACS. The pressure was set at 1bar and controlled with the Berendsen barostat^44^ during initial relaxation; the production runs were carried with the Parrinello-Rahman barostat semi-isotropically with a compressibility of 4.5 x 10^-5^ and a coupling time constant of 5.0 ps.^45, 46^ Non-bonded interactions were computed using a switching function between 1.0 and 1.2 nm, and Particle Mesh Ewald for long-range electrostatics.^47^ The LINCS algorithm^48^ was used to constrain hydrogen bonds in GROMACS.

Simulation parameters for the Anton2 machine were set by ark guesser files, which are automated internal scripts designed to optimize the parameters for the integration algorithms of this machine; as such, the cut-off values to compute interactions between neighboring atoms are set automatically during system preparation. Long-range electrostatics were computed using the Gaussian Split Ewald algorithm,^49^ and hydrogen bonds constrained using the SHAKE algorithm.^50^ Finally, the Nose-Hoover thermostat and MTK barostat were used to control the temperature and pressure during NPT dynamics on Anton2 using optimized parameters set for the *Multigrator* integrator of the machine.^51^

From the MD simulations, we characterized the interactions between protein units as well as with the lipids around them. For the M protein, we also simulated the dimers in water to examine the stability of the proposed structures and to quantify the effect of the bilayer on protein-protein interactions. We computed the root-mean-square-displacement (RMSD) per protein and root-mean-square-fluctuation (RMSF) per residue, and compared it across proposed structures for each protein complex, and versus the proteins in solution in the case of the M dimers. We examined the relative spatial conformations between the proposed models for the M and E complexes, and determined the lipid aggregation patterns around the proteins from micro-second-long simulation trajectories. We carried out one 4-μs long trajectory for each protein-membrane system, and 400ns for the M dimers in water; our reported values are block-averages of the respective quantities. Finally, we compared the degree and extent of membrane deformation caused by each protein complex in terms of the relative position of the lipid headgroups in the cytosolic and luminal leaflets (top and bottom leaflets in our simulation setup). We further discuss below our data in the context of early viral assembly and outline potential directions of this work.

All the images included in this manuscript were rendered on Visual Molecular Dynamics (VMD) software package.^52^ Internal GROMACS modules, and MDAnalysis^53, 54^ and MDTraj^55^ python packages were used to carry the trajectory analysis. Computational time for the trajectories in this work was generously provided by the CODIV-19 HPC Consortium at the Frontera and Anton2 machines.

## Results & Discussion

From studies on other coronaviruses, it is known M and E are implicated in viral assembly, budding, infectivity, and evasion of the immune cell response.^12, 13^ However, little is known about their mechanisms of action, specially their effect on their surrounding lipid environment during viral assembly and in the viral particle itself. First, we report our insights from simulations of an open and a closed M dimer (as seen in Figure 1). Then, we discuss the differences in molecular interactions and membrane response for two proposed models for the E channel. We present a detailed account on the molecular interactions of these integral membrane proteins and comment on plausible mechanisms of action for both proteins in the context of membrane reorganization during early stages of viral assembly of SARS-CoV-2.

The M protein is known as the orchestrator of SARS-CoV-2 viral assembly, preparing it for viral budding and scission. In this context, we examined the interactions between monomers in an M dimer in water and in a lipid environment. Both the open and closed conformations are stable and do not dissociate in water, as shown by the grey and blue structures in Figure 2, and much less so in the bilayer. In fact, the closed conformation barely shifts position due to its tight configuration from the very beginning. The three TMs in the closed conformation form essentially a curved plane, thus have more surface to interact and stabilize the interaction between monomers in the dimer. Similarly, the CTDs of the monomers in this conformation interact very tightly through their smooth faces, as designated in Figure 1. On the other hand, in the open conformation the TMs and CTDs have more flexibility to move. Only TM2 and TM3 interact between monomers, and TM1 is relatively free to move and rotate. The CTDs are separated with their twisted faces point towards the center of the dimer, and the smooth faces point outwards and can easily interact with surrounding lipid headgroups.

**Figure 2.**
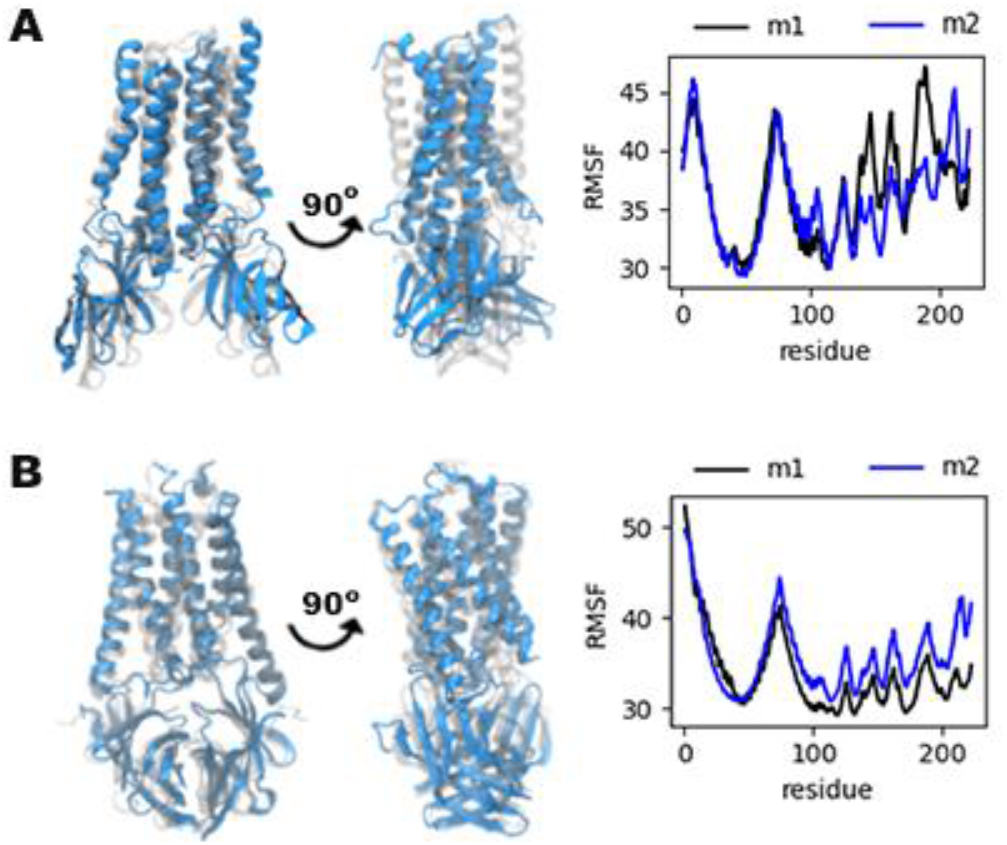
Comparison of the initial (faded) and final (blue) structures of the (A) open and (B) closed conformations of an M dimer in water. Next to each structure is the RMSF computed for the 400ns trajectory of each dimer in water.

Figure 3C&D shows the number of contacts between heavy atoms of each monomer. The dimers in the closed conformation have more contacts than the one in the open conformation, both in water an in a bilayer. As expected from its tighter structure, the TMs in the closed conformation do not fluctuate much in a bilayer environment, and the CTDs come closer together. In fact, the presence of the bilayer restricts the motions of the TMs in the open conformation. Figure 2 shows the TMs in this conformation collapse into each other when in water, bending near mid-region. This is also shown in the accompanying RMSF in the figure, that shows the protein residues in TM1 and TM3 experience large shifts with respect to the initial structure. When the open dimer is embedded into a bilayer, the TMs retain a cylinder-like form throughout the simulation and the TMs of only one monomer experience a large shift from their initial position (see Figure 3.A&C). In the bilayer, the CTDs also fluctuate more in the open conformation versus the closed one, and the CTD from one monomer in the open conformation tilts toward the membrane surface as it interacts with the lipid headgroups.

**Figure 3.**
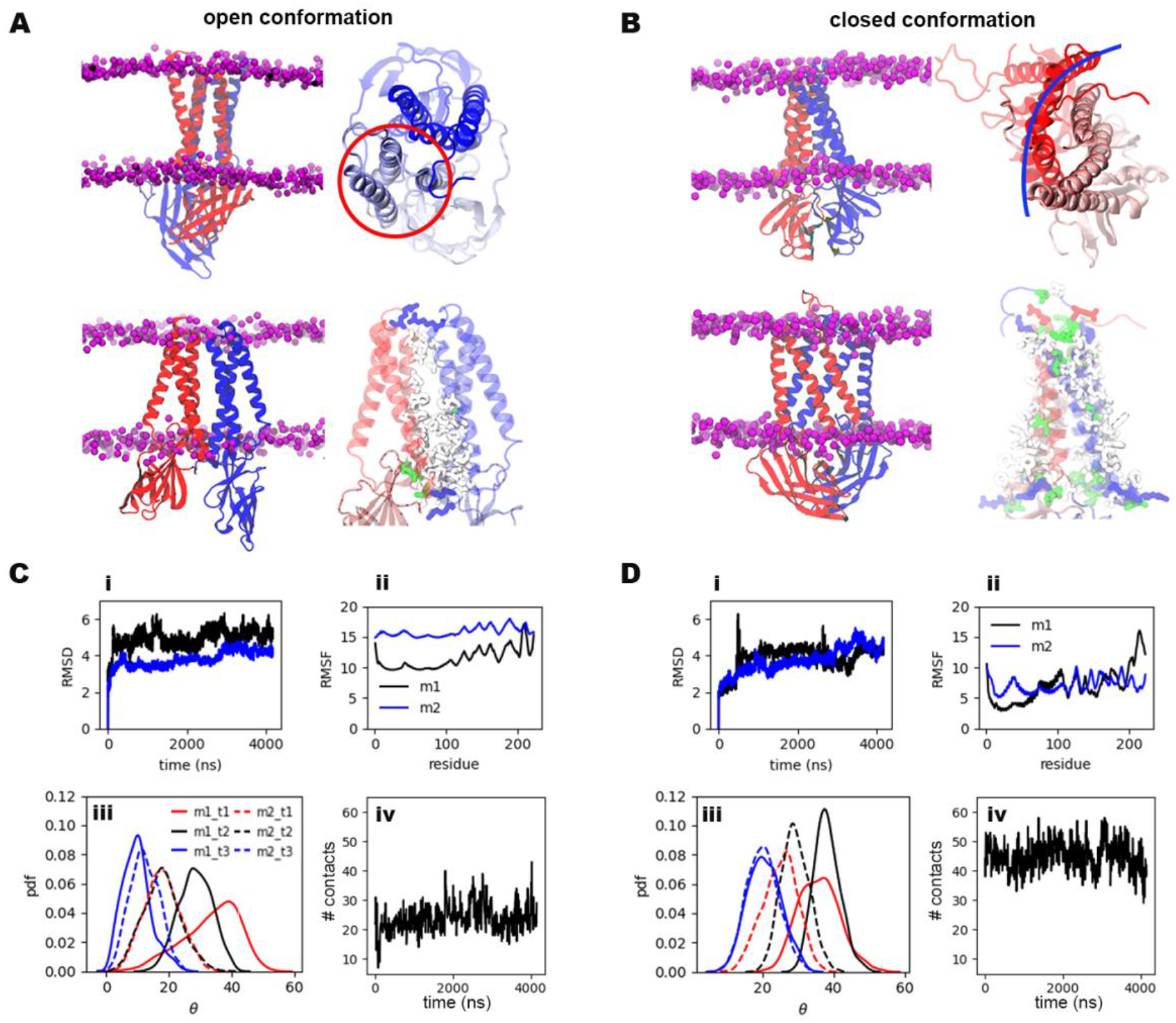
Protein-protein interactions of M dimers in a bilayer. (A) Top and side view of the open dimer conformation highlighting the spatial arrangement of the transmembrane helices and the closest bulky non-polar residues between the protein units. Similarly, (B) the spatial conformation of the closed dimer in a bilayer and non-polar residues pointing towards the bilayer core. Quantitative differences between the (C) open and (D) closed dimers: (i) RMSD of each protein unit. (ii) RMSF per residue per protein unit. (iii) Probability distribution of the angle of each transmembrane helix with respect to the membrane normal. (iv) Number of contacts between M monomers in the respective conformation.

A closer look at the TMs in both conformations inside the bilayer shows that bulky non-polar residues stabilize the interaction between TMs of each monomer in the open conformation, as shown in Figure 3A, whereas these bulky residues point towards the bilayer core in the closed conformation. Looking at the protein motions throughout the simulation trajectory, most of the changes in the open conformation occurred within the first 100ns of simulation (cf. Figure 3Ci), and the displacement of each monomer is not coupled to the movements of the other as shown by the RMSD over the course of the trajectory. On the other hand, the displacement of the monomers in the closed conformation takes longer time and is more coordinated as shown in Figure 3Di. The conformational changes of the closed dimer occur within the first 1000ns of trajectory, then again during the last microsecond. Finally, the orientation of the TMs in terms of their angle with respect to the bilayer normal shows TM1 and TM2 of each monomer align to each other in the closed conformation, and TM3 of one monomer aligns with TM3 from the opposite monomer. In the open conformation, TM1 and TM2 are not aligned in each individual monomer, but TM3 helices align between monomers as in the closed conformation. The angle of TM3 helices in the open conformation is smaller than in the closed conformation, i.e. these helices stand more vertical.

Examining a single M dimer already shows marked differences in the membrane response depending on the dimer conformation. Figure 4 shows lipid density maps averaged over the last 50ns of simulation along with a density map of the proteins in the respective conformation; darker regions indicate higher lipid content. There is more accumulation of cholesterol and inositol (PI) lipids around the open dimer, whereas phosphatidylserine (PS) accumulates near one of the monomers of the open conformation – notably, the monomer whose CTD shifts towards the bilayer. Interestingly, there is larger depletion of DYPE lipids around the open conformation. On the other hand, Figure 4.B shows PS lipids accumulate on both sides of the closed dimer, and DYPE localizes closely around the closed conformation. Charged amino acids at the extremes of the TMs of the closed conformation extend outward from the dimers interface and interact freely with charge lipid headgroups, shown in blue in Figure 3B at the top and the base of each TM. The same amino acids in the open conformation remain mostly trapped between interacting TMs in the dimer (Figure 3A), which can explain the more uneven distribution of charged lipids around the open conformation. Different patterns in lipid sorting also result in different degree of membrane deformation around the dimers.

**Figure 4.**
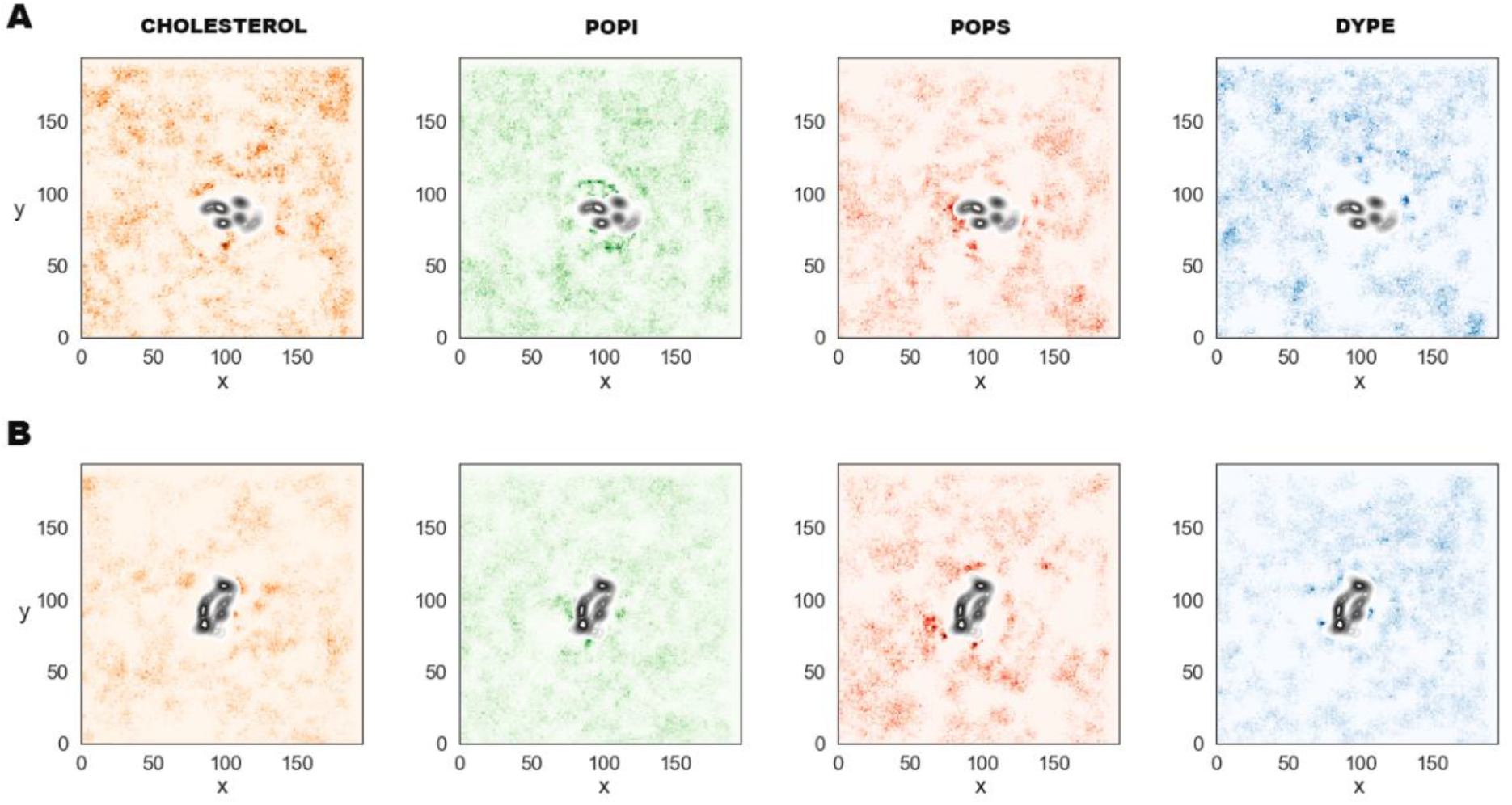
Final lipid density maps relative to the location of the M dimers for the (A) open and (B) closed conformation, Darker regions indicate higher density of the respective lipid species.

Figure 5 shows colored maps of the relative height of each bilayer leaflet in terms of the position of the phosphate groups of each lipid. The red regions correspond to lower positions, and the blue sections are regions where the membrane surface is elevated. A color bar is included per leaflet to clearly distinguish between the lower and elevated regions of each membrane interface. Figure 5A shows the open dimer causes greater deformation on the membrane plane, shifting upwards the lipids around it in both leaflets. From these maps, it is easier to observe the different volume the open dimer occupies across the bilayer. This dimer has a conical shape inside the hydrophobic core of the membrane, with the narrower region in the luminal leaflet and the wider region in the cytosolic leaflet, as observed in the white region at the center of each map. The CTDs of the open dimer are free and able to interact with the lipid headgroups of the luminal leaflet, and push them upwards upon binding. This provides insight into a possible mechanism to generate curvature that contributes to (i) stabilize the budding process of the virus towards the cytosol, (ii) further recruitment of additional viral proteins to the assembly site, and (iii) enhance lipid sorting to modify the mechanical and structural properties of the local membrane in favor of viral particle formation.

**Figure 5.**
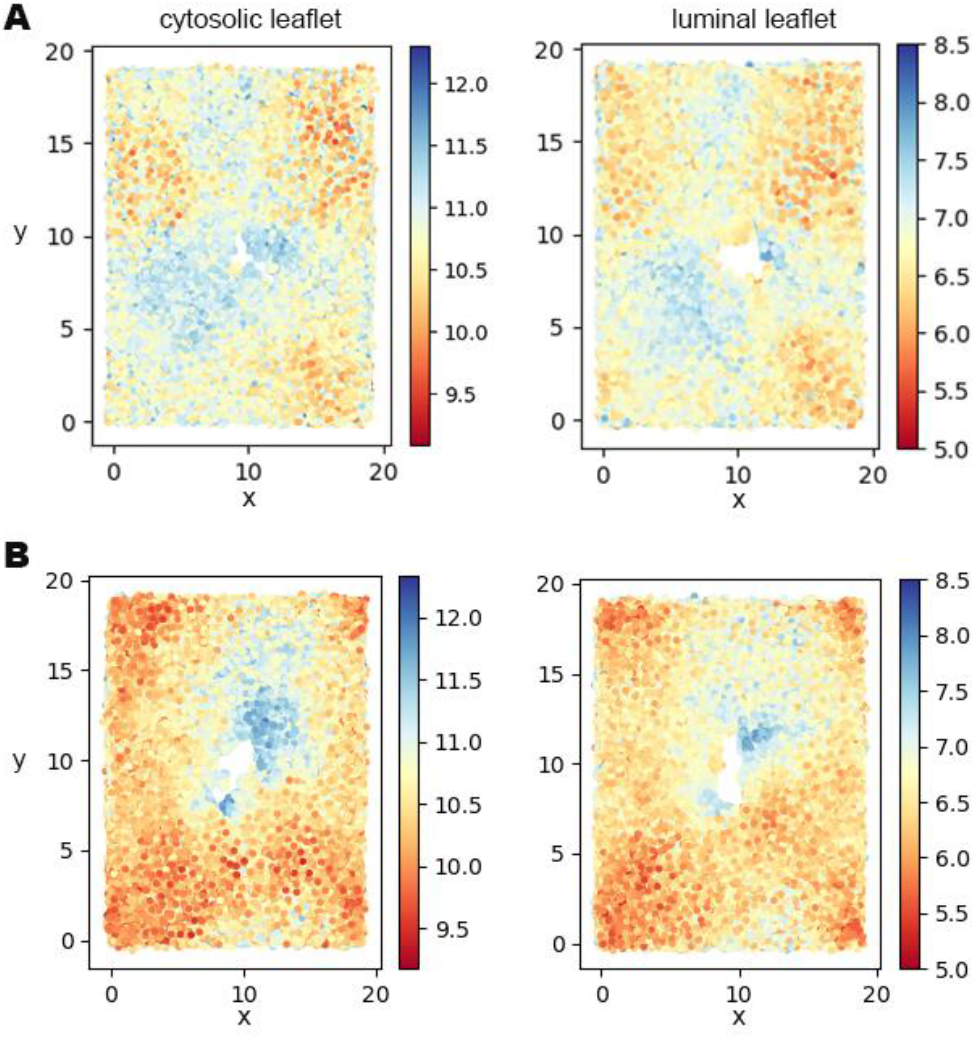
Membrane deformation around the (A) open dimer, and (B) closed dimer. Each map shows the average location of the lipid phosphate groups during the last 50ns of trajectory in the (i) cytosolic and (ii) luminal leaflets, the top and bottom leaflets in the simulation box.

The M protein is the structural protein present in highest concentration in the viral envelope.^9, 12^ Both dimer conformations contribute to membrane deformation towards the cytosol. Given the degree and extent of deformation induced by the open conformation, it is possible this conformation is prevalent at the very beginning to prime the site for viral assembly. As the process advances and the site is more crowded by additional M units and other viral proteins, it is plausible to suggest the open dimers are able to switch partners and adopt the closed conformation, especially since the interfaces of interaction are nearly opposite in both conformations (refer to Figure 1). The closed conformation also pushes surrounding lipids upwards towards the cytosol, but the extent of deformation on the membrane plane is much more local. Figure 5B shows the corresponding lipid elevation maps and an even distribution of the protein inside the bilayer core. In the context of viral assembly, a switch of conformation from open to close dimers and a preference for the closed conformation as the assembly progresses would serve to stabilize a budding virus and further reshape the local lipid landscape to form an optimum assembly platform.

Along with the M protein, the E channel of coronaviruses is also known to actively modulate viral assembly and budding – among other roles.^1, 14^ From our simulations of two proposed conformations for this channel, we observe very different dynamics in the membrane response and the spatial orientation of the channel in the bilayer. We do not observe passage of ions or water in our trajectories for either model, but identified PHE26 in the TM region as potentially involved in the ion function or gating mechanism of the channel. This is the only residue inside the channel that flips its orientation repeatedly during the microsecond-long trajectories. As discussed earlier, PHE26 was also suggested as modulator of the open and closed conformations of the channel in another study^14^ that characterized potential ion channeling mechanisms across homology models similar to the Korkin model used in this work.

Upon examination of Figure 6A, it is clear the Korkin model for the channel causes greater membrane deformation. This occurs in part because the charged residues in the CTDs inside the bilayer core can interact with the lipid headgroups of both leaflets and pull on them like springs. Additionally, as discussed in the next paragraph, the local lipid distribution around the Korkin model is such that it results in a more flexible microenvironment. By contrast, the CTDs in the Feig model remain at the membrane interface and even detach from the surface momentarily to then bind again. The Feig CTDs that detach from the membrane surface can rotate in solvent and bind in a different conformation, which results in the sharp increases in the RMSD profiles of two adjacent monomers in this channel (see Figure 6Aiii) and high fluctuation in the corresponding RMSF profiles at that region. Though the CTDs in the Korkin model move inside the hydrophobic core, the conformational changes are not as pronounced as those in the Feig model as reflected in the corresponding RMSD profiles. Significant conformational changes occur within the first microsecond of simulation in both cases.

**Figure 6.**
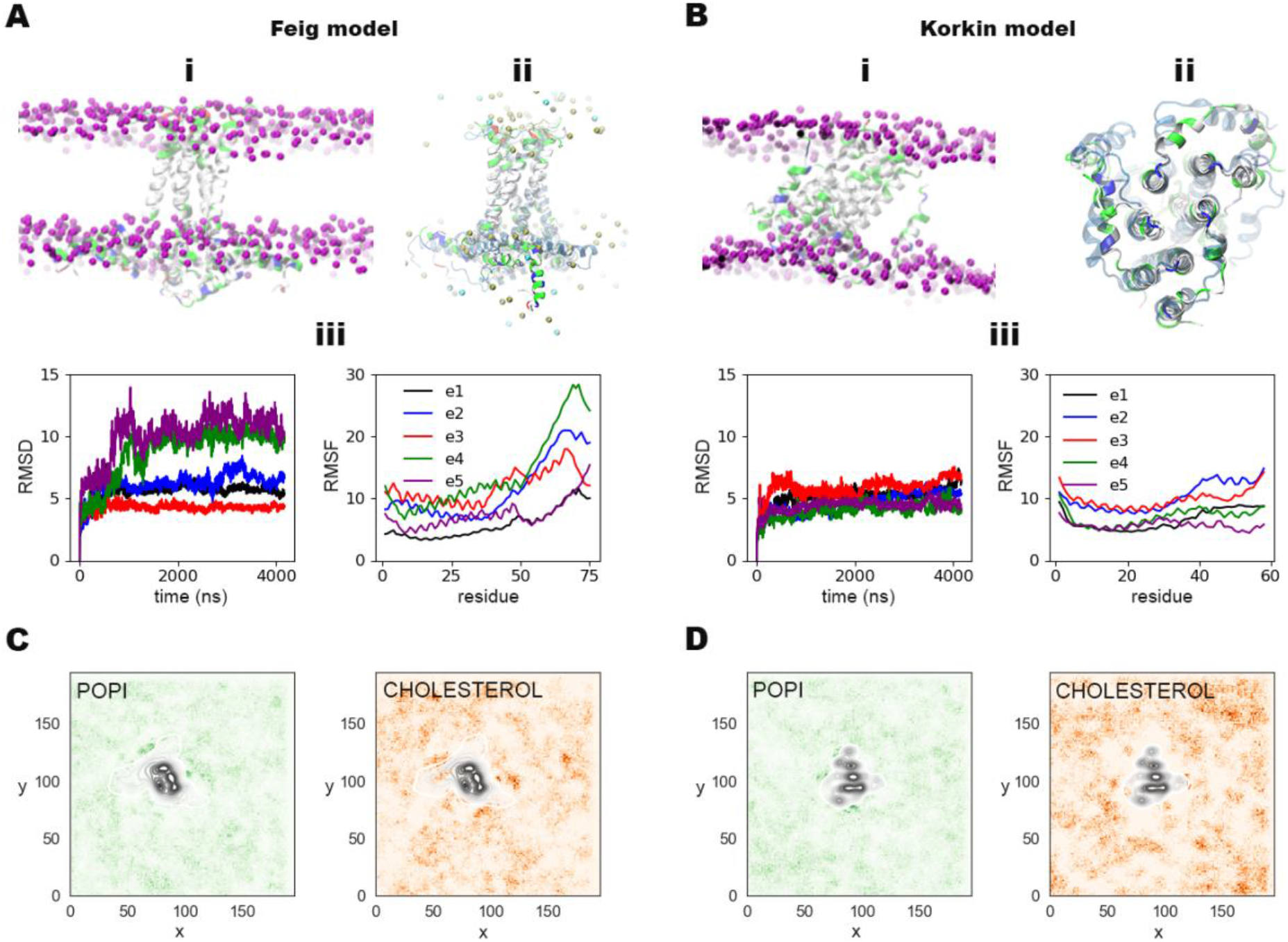
Protein motions of the E channel in a model bilayer. Final snapshots showing the (i) membrane deformation around each channel, (ii) overlaid of the initial (faded) and final channel structures, and the (iii) RMSD and RMSF per protein unit for the (A) Feig and (B) Korkin models; the purple spheres show the relative position of the phosphate group of membrane lipids. (C)-(D) Lipid density plots for inositol and cholesterol lipids around the channel, the protein density is also included in black for reference.

The spatial arrangement of each model induces different lipid rearrangement around the proteins. The hydrophobic TMs of the channel are in direct contact with the bilayer core in the Feig model, whereas the CTDs shield the TMs of the Korkin model inside the bilayer. The CTDs have charged and polar residues that prefer to interact with charged regions of the lipids around them. As a result, the CTDs in the Korkin model attract charged lipid headgroups around them and pull them deeper into the hydrophobic core, shown in darker spots around the protein density in Figure 6C/Di. On the other hand, the TMs in the Feig model, formed by non-polar and bulky residues, attract more cholesterol to the vicinity of the channel (see Figure 6Cii). Cholesterol has a condensing effect, reducing the surface are per lipid and rendering the local environment more ordered and rigid. The opposite occurs around the Korkin model, which has a cholesterol-depleted region around the protein, where the local environment is more flexible. The channel itself, i.e., the relative position of the TMs with respect to each other, does not fluctuate much as shown in the overlaid snapshots in Figure 6A/B, but there are pronounced shifts of the CTD in both models as these actively interact and sort the lipids around the protein.

The difference in local environment around each model allows or restricts further protein motions. There are three phenylalanine rings in the TMs of the channel: PHE20, PHE23 and PHE26; only the last one is completely inside the channel during the entire trajectory and is also suggested as key modulator between the open and closed conformations of the channel.^14^ Figure 7A-B show 2D histograms for PHE26 residues in the TM of each model (the equivalent residue in the Korkin model is PHE19, as listed in Figure 1C). The vertical axis corresponds to the rotation of the PHE ring plane, denoted by the cosine of the angle between the vector formed by the adjacent carbons to the ring tip (CE1 and CE2 in the protein topology file) and the bilayer normal; a value of one indicates the PHE ring plane is perfectly aligned to the bilayer normal or z-axis in our simulation systems, where as a value of zero indicates the ring plane is parallel to the bilayer center plane. The horizontal axis corresponds to the PHE flip inside the channel, denoted by the cosine of the angle between the vector formed by the base and tip carbons of the ring (C_β_ and C_Z_ in the topology) and the bilayer normal; a value higher than 0.5 indicates PHE points upwards in our simulation coordinates, towards the luminal side or the interior of the virion, and values lower than 0.5 indicates the PHE ring is nearly flat aligned with the bilayer center plane. The PHE residues in the TM in the Korkin model have higher incidence, in number of TMs, and frequency, in number of flips, of orientation changes. Even the probability density profiles show two distinct peaks for PHE rings in the TMs in the Korkin model (Figure 7D).

**Figure 7.**
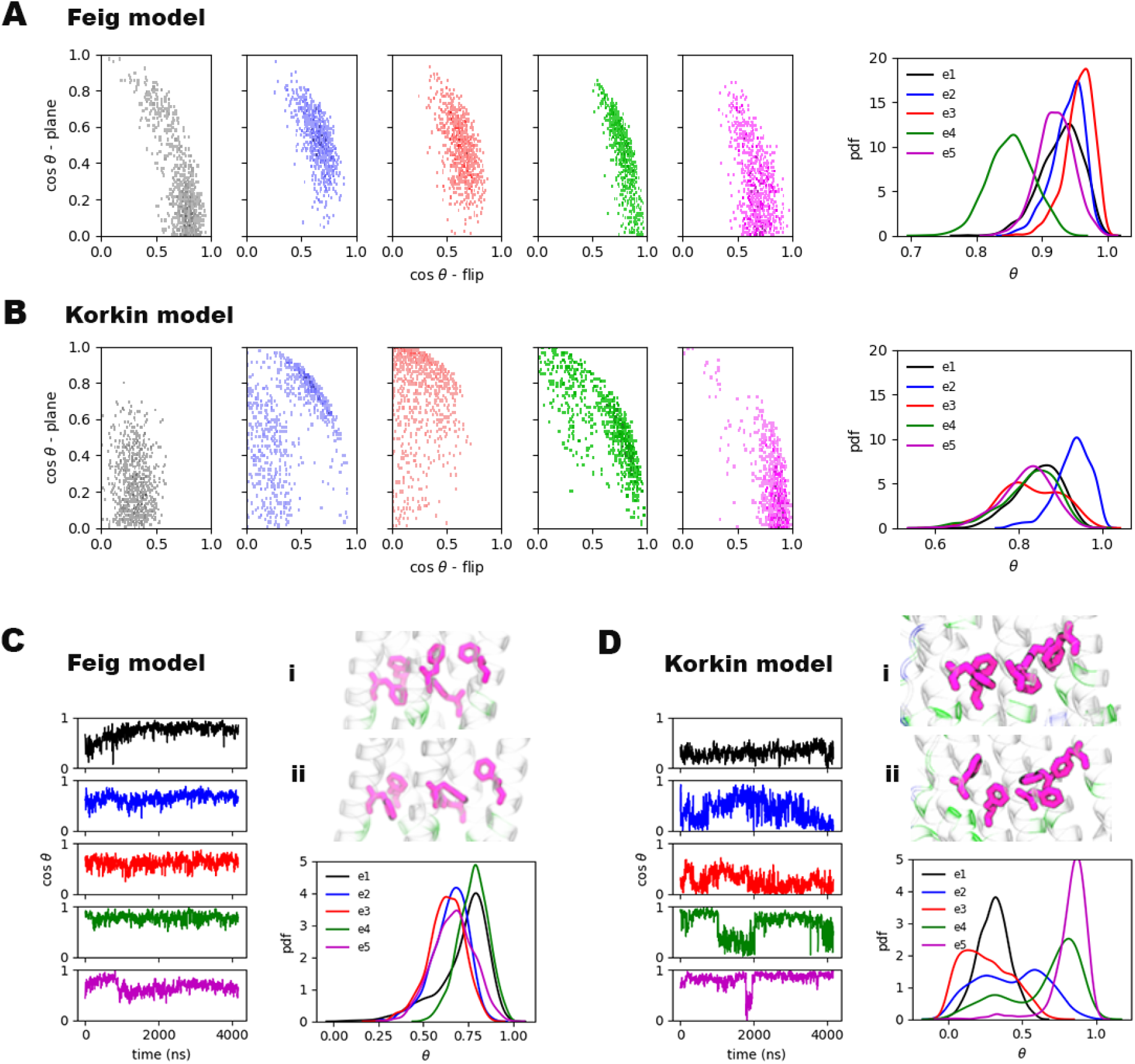
Rotation maps to quantify the motions of the PHE ring plane and it rotates and flips inside the channel for the (A) Feig and (B) Korkin models, and the corresponding probability distribution of the TMs wrt. the bilayer normal in each model. (C)-(D) Time series and probability distribution of the cosine of the PHE26 angle wrt. bilayer during the trajectory for the Feig and Korkin models, respectively; also shown are the (i) initial and (ii) final orientations of PHE26 inside each channel.

We observed a more symmetric response in the membrane leaflets surrounding the Feig model in terms of the position of the P atoms. Though there is a change in the topology of the membrane surface from the initial to the final set of frames in the simulation, the change is coordinated between the leaflets, the same regions in the xy-plane are elevated or depressed. As shown in Figure 8, the bilayer is pulled towards the cytosolic leaflet at the region where the CTDs of the Feig model are in contact with the membrane surface (shown in red in panel 8A). On the other hand, the leaflets surrounding the Korkin channel vary quite differently; the cytosolic leaflet remains pretty much at lower values, there is barely a small elevation in blue right at the edge of the pore created by the protein. Whereas the luminal leaflet, initially at a uniform elevation, reshapes to nearly half of its surface at 1.5-2 angstroms higher than the other half. Due to the CTDs inside the bilayer core, the cross-sectional area of the Korkin model is much more irregular than the highly conserved shape of the Feig model.

**Figure 8.**
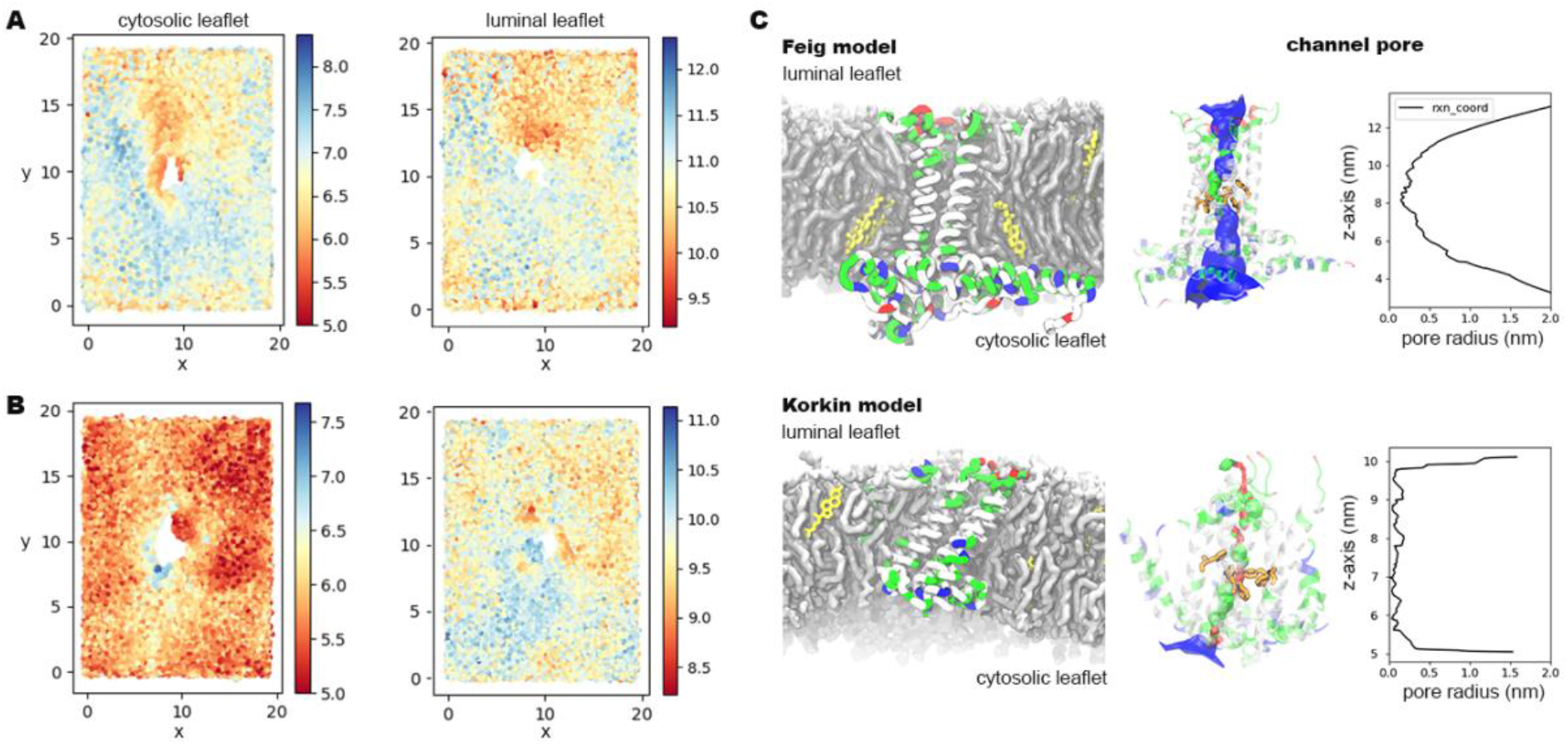
Membrane deformation around the E channel: (A) Feig and (B) Korkin models. Each map shows the average location of the lipid phosphate groups during the last 50ns of trajectory in the (i) cytosolic and (ii) luminal leaflets, the top and bottom leaflets in the simulation box. (C) The Feig and Korkin models for the E channel shown as reference in the bilayer with a corresponding snapshot of the channel pore at the end of simulation computed on MDAnalysis implementing the HOLE^56^ routine; the blue regions can accommodate two water molecules, green regions have enough space for one water molecule, and red regions are too narrow to allow the passage of a water molecule.

It is clear that the protein-lipid interplay around the E-channel is very complex. An additional homology model^14^ has been proposed since the start of our study, simulations of an E monomer to characterize the orientation of the helices in the sequence,^57^ an experimental structure for the TMs (7K3G).^11^ It remains to be determined if the preferred E channel conformation resembles the Korkin model, or if a conformation like the one proposed by the Feig model is biologically relevant at a given stage of the viral life cycle. From previous studies, it seems the Korkin-like conformation is prevalent when the E channel functions as an ion channel, yet the channeling activity is highly modulated by local lipid environment.^14, 15^ Our results also favor that premise in the Korkin model could presumably let ions pass through with some selectivity, based on the size of the channel pore identified and shown in Figure 8C. The Feig model, instead, remains wide enough throughout the 4μs trajectory to accommodate up to two water molecules in its widest regions, identified by the blue coloring of the channel pore. Latest studies on the function of the E channel suggest it allows the passage of both monovalent and divalent cations modulated by the local lipid composition.^15^ Furthermore, simulation studies suggest the glycosylation of the CTD helices could also determine their orientation with respect to the lipid bilayer.^57^ There is a narrow area right at the location of PHE26, which our results also point as potentially key modulator for the ion channeling activity of the complex.

Since both E and M are implicated in viral assembly and budding, further modeling and simulation studies that examine their coordinated dynamics in the bilayer as well as in higher individual concentration are needed. Enhanced free energy sampling methods that enable us to quantify the energetics of protein-protein (M-M) and intra-protein (M-E) interactions would also serve to further understand the function of these integral membrane proteins in the assembly and budding of SARS-CoV-2 and related viruses. All-atom MD simulations offer an advantage to understand specific protein-lipid interactions and motions; however, a more multiscale approach is likely needed to gain better understanding of this intricate system.^58^

## Conclusions

This work summarizes our initial efforts to understand specific protein-lipid interactions of two structural proteins of SARS-CoV-2. Specifically, we examined differences between two proposed homology models for a dimer of the M protein and the viroporin channel formed by five monomers of the E protein. We used all-atom MD and a complex membrane model to determine protein motions and interactions in a lipid environment that closely resembles the place where viral particles co-localize during early stages of viral assembly, the ERGIC. We propose that both of the two M dimer conformations suggested from homology models are present during viral assembly, potentially in different ratios depending on the stage of assembly and maturation. We discuss the effect of both the M and E viral proteins on lipid-lipid interactions and sorting around the proteins, and the potential impact of these interactions on membrane deformation during viral assembly. Finally, we identified PHE26, a residue located on the inside of the TMs of the E channel, as possibly involved in the ion gating function of this protein complex. Data from experiments also identified PHE26 pointing towards the E channel, inside the pore.^11^ PHE 26 is one of the three PHE residues in the TM region of the channel that greatly shifts in the Korkin model. From experimental observations, all three residues are also identified as relevant modulators of the channel’s function along with local lipid composition, which largely modulates the channel’s gaiting activity.^11, 15^ Taken together, these observations seem to identify the Korkin model as the prevalent conformation when E serves as an ion channel, and the Feig model as a potential alternate conformation during other functions of the E viroporin. All the protein conformations we have examined in this work have a different effect on the immediate lipid environment, which is a key feature during viral assembly and membrane remodeling.

We continue to refine our simulation models in collaboration with experimental partners to provide further insights into the complex relationship between viral proteins and lipids in the cell. In addition to providing an atomistic view of these molecular interactions, data from this work can be used to systematically refine our multiscale coarse-grained model of the SARS-CoV-2 virion to better understand the collective dynamics of the entire viral particle.^58^ Our simulation framework also enables us to update our models as new structures and insights from experiment become available. There is certainly pressing need to obtain experimental structures of the M and E proteins that will allow us to examine more closely their interactions and impact on lipid and viral dynamics in the cell. Gaining clear understanding of the viral mechanisms of SARS-CoV-2 is furthermore critical to the development of novel treatments, repurpose existing ones more efficiently, and anticipate strategies to combat natural mutations of this and related viruses.

## Author Contributions

G.A.V provided funding, resources, and supervision. V. M. worked on conceptualization, data curation, formal analysis, and writing of the initial draft. G.A.V and V.M. reviewed and edited the final manuscript. The authors thank the Voth Group, especially Dr. Alex Pak, for insightful discussions on the simulation systems at the beginning of this work. We also thank Prof. Rommie Amaro and her team for initial discussions about interactions among structural membrane proteins of the virus.

## Acknowledgements

This work was supported in part by the National Science Foundation through NSF RAPID grant CHE-2029092 and in part by the National Institute of General Medical Sciences of the National Institutes of Health through grant R01 GM063796. Computational resources were provided by the COVID-19 HPC Consortium (project MCB20037); specifically, Frontera at the Texas Advanced Computer Center (TACC) and the Anton 2 machine at the Pittsburgh Super Computing Center (PSC) (available under NIH Grant R01GM116961). The Anton 2 machine at PSC was generously made available by D.E. Shaw Research. Initial equilibration of some of the systems in this study was computed on Midway2 at the Research Computing Center (RCC) at the University of Chicago.

